# Prediction of binding sites of GPCRs based on 3D convolutional neural networks

**DOI:** 10.1101/2024.06.11.598048

**Authors:** Junfeng Yu, Ying Zhang, Jun Lv

**Author notes:** Corresponding authors (Y.Z.) and (J.L.).

## Abstract

G protein-coupled receptors are a class of receptor proteins located on the cell membrane, regulating the perception and response of cells to various external signals. Identifying the binding sites of G protein-coupled receptors plays a crucial role in understanding their allosteric modulation mechanisms. However, obtaining the crystal structure of the complex through experimental means and subsequently identifying the binding sites require substantial resources. With the development of computer-aided computation, deep learning can effectively predict the binding sites between proteins and ligands. This study predicted the binding sites of G protein-coupled receptors based on 3D convolutional neural network. A total of 108 G protein-coupled receptors recorded in the scPDB database were collected for this study, and a 3D convolutional neural network model was established based on these three-dimensional structures. Firstly, the PDB file of the protein is voxelized and segmented into different channels according to the type of atoms in a certain region. Then, 3D convolutional neural network is employed to predict the binding site through traversal, and the optimal voxel box size and model parameters are determined based on performance evaluation metrics. The established 3D convolutional neural network accurately predicts the binding site of G protein-coupled receptors, with an accuracy as high as 0.942, precision of 0.678, and recall rate of 0.532. Additionally, using the backpropagation algorithm, the gradients of the input data are calculated, and the importance of input elements on the final classification result is analyzed.

## 1 Introduction

G protein-coupled receptors (GPCRs) are a class of proteins located on the cell membrane. They transmit signals inside and outside the cell and regulate a variety of physiological processes. GPCRs work by changing their shape and switching between different conformations to bind with external molecules, which triggers signaling pathways and initiates specific physiological responses (Latorraca et al., 2017). Due to their importance in physiological and pathological processes, GPCRs have become one of the important targets in drug development. Identifying the functional sites of GPCRs to design drugs to treat various diseases is of significant importance (Cheng et al., 2023). Identifying the potential binding sites of GPCRs is one of the important research fields in bioinformatics. Although the binding sites can be identified by obtaining the crystal structure of complexes through experimental crystallization, this method generally consumes a large amount of resources and cannot predict some potential functional sites (Katritch et al., 2013). If the binding sites of proteins could be effectively predicted in advance with the help of computer-aided calculations before experimentation, it could reduce experimental costs and facilitate the design of ligands (Broomhead et al., 2017). With the development of computational science, effective predictions of protein-ligand binding sites have been made through computational approaches such as molecular docking, molecular dynamics simulations, and machine learning, overcoming the shortcomings of traditional experimental methods (Zhao et al., 2020).

The biological function of proteins depends on their three-dimensional structure and specific conformational changes. Molecular docking searches for potential binding sites and conformations in the structure of proteins and ligands, typically using algorithms and energy scoring to measure binding affinity, thereby assessing their interaction capabilities (Li et al., 2019). Molecular docking is mainly used to predict the binding mode between proteins and small molecules, that is, to identify the binding sites of the protein as well as the optimal binding conformation of the ligand (Ferreira et al., 2015). Natasja et al. (2003) focused on developments related to the field of molecular docking, analyzing the forces that play an important role in molecular recognition and interpreting these forces with the help of different scoring functions. Thomsen et al. (2006) developed the MolDock docking algorithm, which combines differential evolution with cavity prediction algorithms and introduces a new rescoring function, able to identify the correct binding modes of 87% of the complexes. Shoichet et al. (2012) briefly introduced the current status of computational molecular docking on GPCRs, pointing out that the binding pockets of GPCRs are highly suitable for docking and can be used to screen for effective and novel compounds for targets. Ballante et al. (2021) reviewed the widespread application of molecular docking in GPCRs-ligand complex modeling and chemical probe screening, noting that structure-based virtual screening is key to discovering new GPCRs drugs. Muthiah et al. (2021) used existing anticancer drugs as ligands for molecular docking with GPR116, carefully analyzing the molecular interactions to identify key residues in the binding pocket.

Molecular docking focuses on the static interactions between molecules, while molecular dynamics simulations pay attention to the motion and dynamic processes of biomolecules. Molecular dynamics simulations reveal protein structure and function by tracking protein conformational changes, such as protein folding, ligand binding, and enzymatic catalysis, providing important insights into biological phenomena (Liu et al., 2018; Re et al., 2011). Ichiye et al. (1991) studied the covariance of atomic fluctuations in molecular dynamics and normal mode simulations to identify collective motions within proteins, comparing the covariance matrices and cross-correlation matrices to find many similar features of relative motion in different regions of the protein. Borhani et al. (2012) explored new directions for future drug research based on molecular dynamics simulations, recognizing the great potential for the development of molecular dynamics simulations. Sengupta et al. (2015) used molecular dynamics simulations to successfully explore the effects of cholesterol on receptors, emphasizing the main characteristics of interactions between cholesterol and GPCRs, and identifying the binding sites on receptors that interact with cholesterol. Although molecular dynamics simulations can accurately predict protein binding sites, they usually involve significant expense. Simulating large protein conformational changes, such as those in GPCRs, is challenging to study with molecular dynamics simulations. Brooks et al. (1983) proposed a coarse-grained normal mode analysis (NMA) to overcome the time-consuming and high-cost difficulties of molecular dynamics simulations, and successfully analyzed useful results of internal motions in bovine pancreatic trypsin inhibitor. Tirion (1996) optimized and improved NMA by proposing the elastic network model (ENM), which more effectively simulates protein functional motions. By calculating the root-mean-square fluctuations of different modes, this model accurately predicted the structural flexibility of proteins.

In recent years, based on the development of machine learning methods, machine learning began to be used to predict the binding sites of protein. These methods can be roughly divided into sequence information and structure information based on protein (Marrone et al., 1997). The sequence-based method uses the evolutionary conservation or sequence similarity from homologous protein and the specific characteristic information of amino acids, while the structure-based method uses the three-dimensional atomic coordinate information of protein, then extracts feature engineering, uses machine learning algorithm to select features, eliminates redundant or irrelevant features, improves the generalization ability of the model, and finally infers potential binding sites and key residues (Zhao et al., 2020). In terms of protein feature engineering, features such as the position-specific scoring matrix (PSSM), sequence one-hot encoding, amino acid physicochemical properties, secondary structure, functional and structural domains, solvent accessible surface area, and distances between residues are commonly used. Lin et al. (2005) used biofeature encoding technology and established a neural network using a sliding window technique to predict protein metal binding sites. Capra et al. (2009) combined evolutionary sequence conservation information with other structure-based methods, advancing the understanding of the relationship between evolutionary sequence conservation and protein structure and functional attributes. Nikam et al. (2023) established the deep neural network DeepBSRPred to predict protein binding sites based on specific sequence and structural features, using parameters such as the position-specific scoring matrix, solvent accessible surface area, conservatism scores, and amino acid properties. Le et al. (2019) created position scoring matrix maps and used two-dimensional convolutional neural networks to accurately identify GTP binding sites in Rab proteins. Jiménez et al. (2017) used binding site information from the scPDB database, calculated atom occupancy in proteins based on atomic characteristics, assigned atoms to specific channels, and created a 3D convolutional neural network to predict protein binding sites.

This paper establishes a 3D convolutional neural network based on the three-dimensional structure of proteins to recognize the binding sites of G-protein coupled receptors. The three-dimensional structural data is mapped and dimensionally reduced to generate two-dimensional planar data, and the 2D convolutional neural network is used for prediction, which is compared with the predictions of the 3D CNN. At the same time, machine learning models like SVM are used to predict and analyze the one-dimensional data of the structure. The model’s performance after training is evaluated using different index systems. This study is the first to use a 3D convolutional neural network to predict and analyze the binding sites of GPCRs, which is instructive for the design of related receptor drugs.

## 2 Data Processing and Model Building

### 2.1 Data collection and processing

This article uses PDB data provided by the GPCRdb (https://gpcrdb.org/) database as the raw data. This database collects structural and sequence information for 1160 GPCRs (Munk et al., 2016). In order to study the binding sites of GPCRs, the protein-ligand binding site data recorded in the scPDB database is used as the target function. The scPDB database (http://bioinfopharma.u-strasbg.fr/scPDB/) is a structural database focused on research into protein and ligand binding sites, and it records the binding site data for 17594 proteins (Desaphy et al., 2015). This database includes information on protein structures, ligand molecules, protein cavities, and binding patterns. By filtering the binding site information of 108 GPCRs recorded in the scPDB database, we established our GS108 dataset. This dataset includes multiple families of G protein-coupled receptors, with a total of 40620 amino acids. We use 25 different proteins of A_2A_AR as the test set, containing 8752 amino acids, and the remaining 83 proteins as the training set. Based on the binding site information recorded by scPDB, the amino acids in the dataset are one-hot encoded. If the residue of the protein is a binding site, it is marked as a positive sample with a 1; otherwise, it is marked as a negative sample with a 0. Detailed information of the dataset is shown in Table 1.

**Table 1.** Composition of training set and test set.

| Data set | Sequence number | Positive sample | Negative sample | Sparsity rate |
| --- | --- | --- | --- | --- |
| Training set | 31868 | 2874 | 28994 | 9.91% |
| Test set | 8752 | 699 | 8053 | 8.68% |
| Total | 40620 | 3573 | 37047 | 9.64% |

Note: Sparsity rate is the ratio of negative sample number to positive sample number.

Since the input data of the 3D convolutional neural network requires 3D multi-channel, it is necessary to voxelize the three-dimensional structure of the protein. Voxelization encodes the spatial arrangement of atoms and atom types into input data for the model so that individual atoms are also accurately characterized. The voxelized three-dimensional space not only retains the three-dimensional atomic structure information, but also provides an ideal data input for CNN (Bouvier, 2021). We only have the coordinate file data of the protein, and for a given amino acid, we use the Cα atom as the center and traverse to search for other atoms within a certain distance, so that the relevant atoms can be voxelized by the coordinate position alone.

In order to standardize the data, coordinate transformation is carried out for each atom, with the vector from Cα atom to N atom as the X axis and the vector from Cα atom to another C atom as the Y axis, and then the Z axis is calculated by cross product. Finally, the direction of Z axis is adjusted by calculating the direction between Cβ and Cα atoms, so as to maintain the original positional relationship between atoms as much as possible. In this paper, a voxelized space with dimensions of 32 × 32 × 32 (Å^3^) is established with the Cα atom of the amino acid as the center, and then the space is sliced in units of 1 Å, and each voxel is a cube with a side length of 1 Å. Atom type channels were used to record the states of existence of carbon, oxygen, nitrogen, and sulfur atoms, and the corresponding atoms in the amino acids were separated to produce 4 × 32 × 32 × 32 dimensional list data. Splitting the space in 1Å units ensures that each voxel holds only one atom, thus achieving better spatial resolution. The initial value of all voxels is 0. If the Cartesian coordinates of an atom of a certain type are within a voxel, then the voxel of the corresponding channel is assigned the value 1. Because the voxels recording atomic information are too sparse, Gaussian smoothing method will be used to diffuse and fill the structure list data, and the original structure data will be enhanced, as shown in Figure 1.

**Figure 1.**
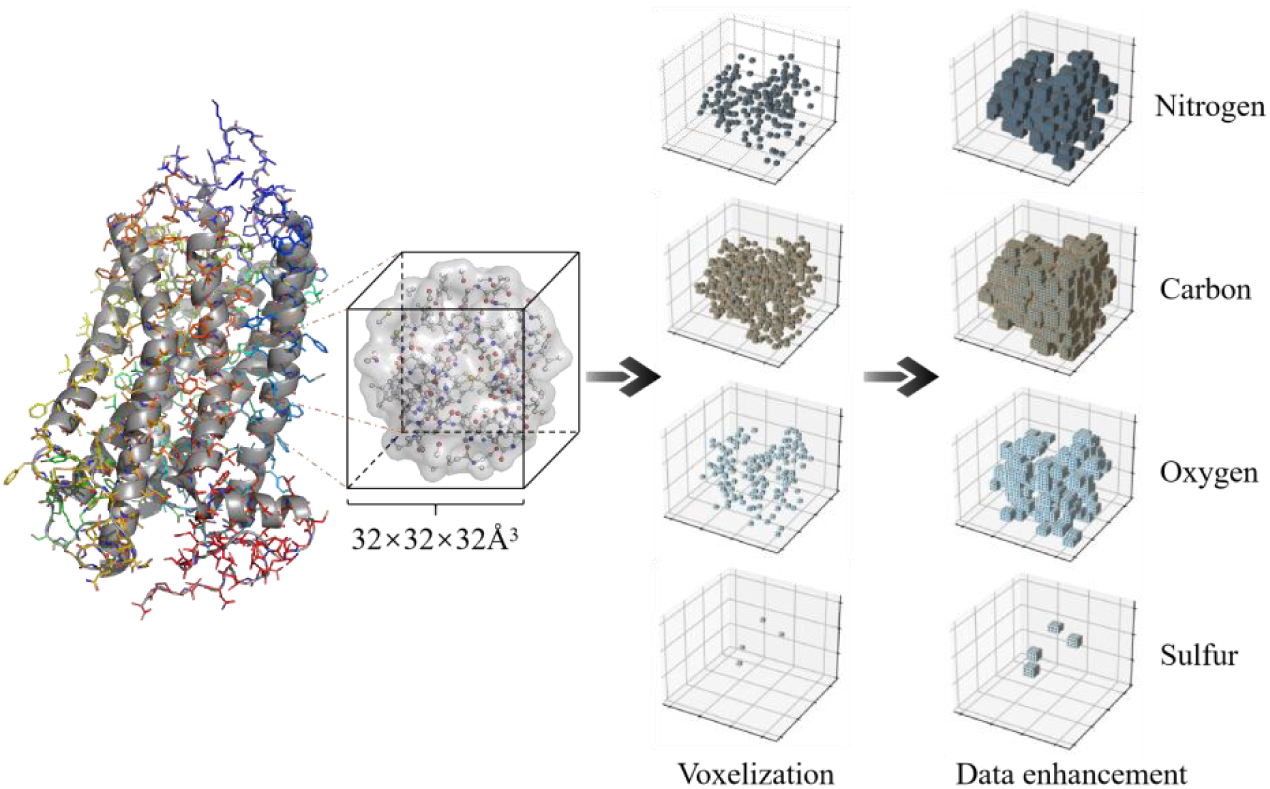
Divide channels according to atomic types for voxelization.

### 2.2 Constructing 3D convolutional neural network

3D Convolutional Neural Network (3D CNN) is a type of deep learning model used for processing three-dimensional data. Compared to traditional two-dimensional convolutional neural networks (2D CNN), 3D CNN are better at capturing spatio-temporal information in three-dimensional data such as videos and volumetric structures. The structure of 3D CNN is basically the same as that of 2D CNN, mainly composed of convolutional layers, pooling layers, fully connected layers, etc. However, the difference is that the convolutional kernels in a 3D CNN are three-dimensional, allowing them to slide across three dimensions simultaneously, thus effectively capturing spatio-temporal features. Machine learning typically requires physical and chemical features as input, whereas the 3D CNN method automatically extracts features from GPCRs protein structures, achieving prediction of binding sites with less information (Jiménez et al., 2017). Figure 2 shows the structure of the three-dimensional convolutional neural network constructed in this chapter and a specific parameter flowchart.

**Figure 2.**
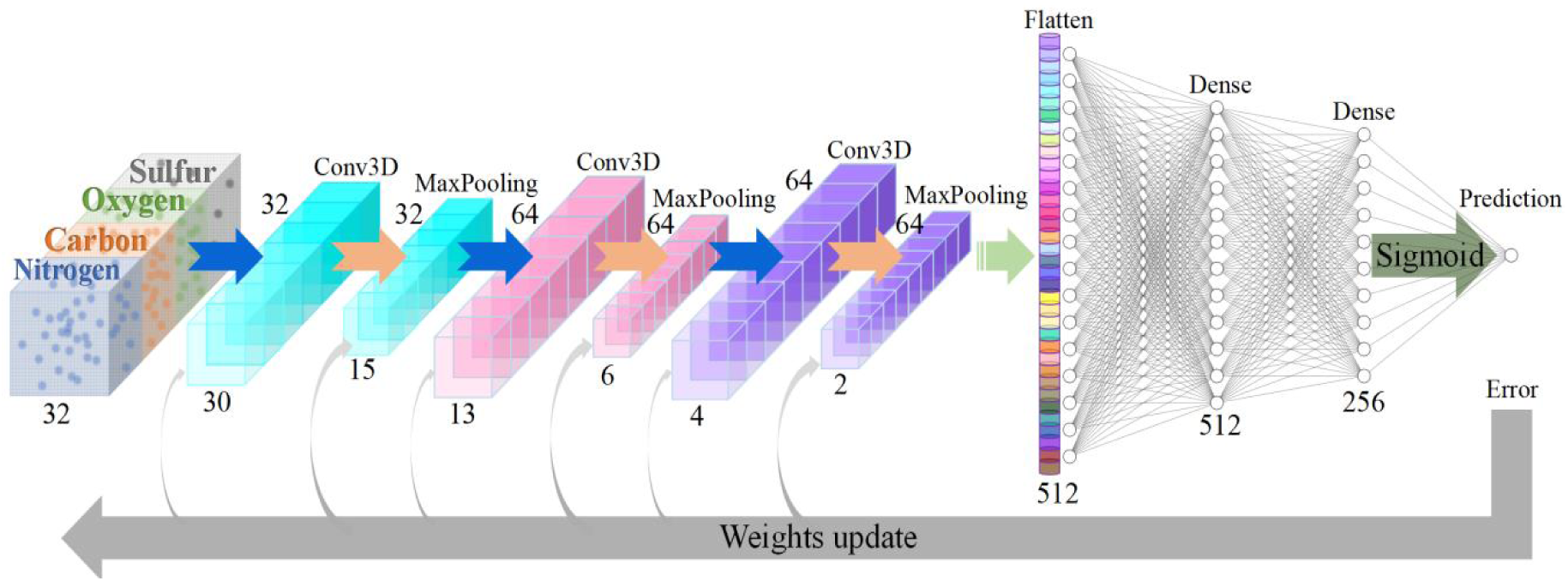
Flow diagram of 3D convolutional neural network model.

This article builds a Convolutional Neural Network architecture based on the Keras framework, using the three-dimensional structure of GPCRs to classify and determine the binding sites. The voxelized GPCRs structure is input into a model that contains a three-layer 3D CNN architecture for feature extraction, and the convolutional layer extracts the spatial features of the structure in three dimensions. The first convolutional layer (Conv3D) has 32 filters, with a kernel size of (3,3,3), and uses the ReLU activation function, with the weights randomly initialized using kernel_initializer=’he_uniform’. The second convolutional layer has 64 filters, and the third convolutional layer also has 64 filters, all using the same kernel size and activation function. After each of the three convolutional layers, a max pooling layer has been added with a pool size of (2,2,2). A dropout layer with a dropout rate of 0.2 has been added after each convolutional layer in the model. The dropout layer regularizes the network to prevent overfitting. Then, a Flatten layer is added to convert the 3D feature map into a 1D tensor, for input into the fully connected layer. The network contains two fully connected Dense layers, with 512 and 256 neurons respectively, using ReLU activation and he_uniform initialization. The final output layer is a Dense layer with a single neuron and a sigmoid activation function, which is used for binary classification. The convolutional network we designed uses the feature tensor to predict the binding sites of GPCRs. Figure 3 shows the workflow of this chapter.

**Figure 3.**
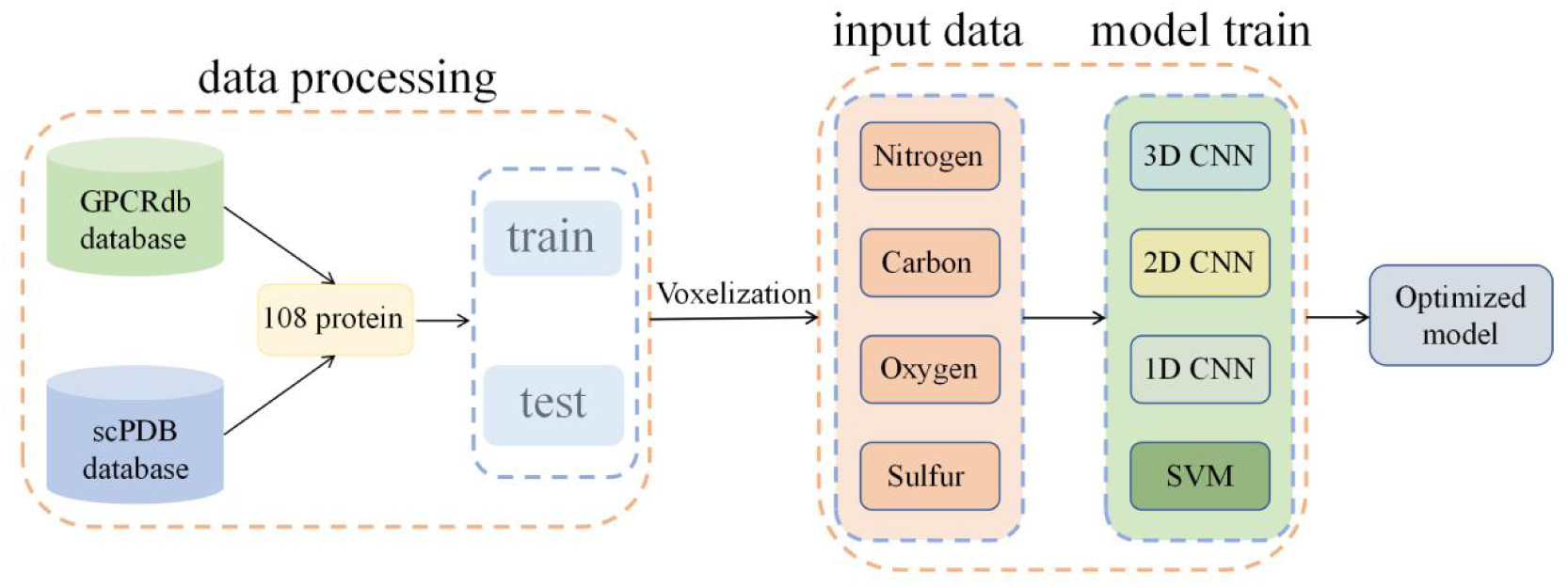
Schematic diagram of workflow, including data acquisition and reconstruction workflow, model training and inspection.

### 2.3 Model evaluation

The confusion matrix is an important tool in machine learning and statistics for evaluating the performance of classification models. It shows the correspondence between the prediction results of a classification model on a test dataset and the actual labels. TP represents the number of positive samples correctly predicted as positive by the model, TN represents the number of negative samples correctly predicted as negative by the model, FP represents the number of negative samples incorrectly predicted as positive by the model, and FN represents the number of positive samples incorrectly predicted as negative by the model. Through the confusion matrix output by the model, we can understand the accuracy and reliability of the convolutional neural network predictions, and better evaluate the performance of the classification model. The confusion matrix provides the basis for calculating a number of important classification metrics, including but not limited to Accuracy, Specificity, Recall, Precision, Matthews Correlation Coefficient (MCC), and F1-score. These metrics can comprehensively and accurately evaluate the performance of the classification model, thereby helping us choose the most suitable classifier for the task. Therefore, the confusion matrix becomes one of the indispensable tools for evaluating the performance of classification models.

(1) Accuracy: The sum of the true positive and true negative samples predicted by the model in the dataset, compared to the total number of samples in the test dataset. It is a common metric for evaluating the performance of a classification model, usually expressed as a percentage.

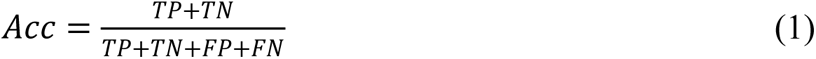
(2) Specificity: The ratio of the true negative samples predicted by the model in the dataset to the total number of actual negative samples, that is, the proportion of actual negatives that are correctly identified as such.

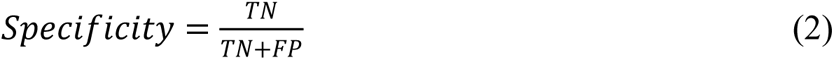
(3) Recall, also known as sensitivity or hit rate, measures the proportion of actual positive samples that are correctly identified by the classifier out of all actual positives. A high recall usually implies a low misclassification rate and high accuracy.

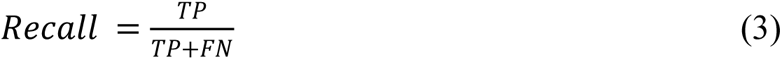
(4) Precision, also known as positive predictive value, is a crucial metric for evaluating the performance of binary classification models. It measures the accuracy of the model when predicting positive outcomes and is often used in conjunction with recall and other metrics to provide a comprehensive and precise evaluation of a classification model’s performance. A higher precision indicates better effectiveness of the model. In some scenarios, such as dealing with high-volume data or imbalanced class distributions, precision can be especially important.

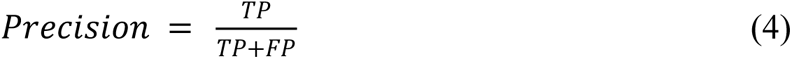
(5) The Matthews Correlation Coefficient (MCC) is a measure used for assessing the performance of a classification model. A value closer to 1 indicates a more perfect prediction outcome for the test subjects.

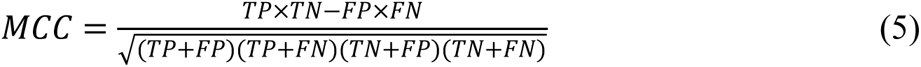
(6) The F1-score is the harmonic mean of Precision and Recall. It is used to measure the performance of a classification model, especially in instances of imbalanced class distribution. It can be calculated by the following formula:

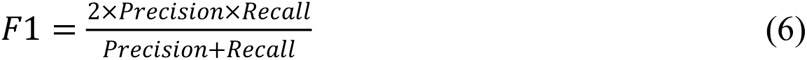

To evaluate the merits of a prediction model, it’s often necessary to present a comprehensive view of the various metrics since each metric focuses on a specific aspect, and their performance is usually linked to the selection of the classification threshold. Thus, citing a single metric alone cannot objectively reflect the performance strengths or weaknesses of a classification model. In this article, the AUC (Area Under the Curve) will be calculated to assess the performance of a 3D CNN model. The AUC value ranges from 0 to 1, where 0.5 signifies that the model performs no better than random guessing, and 1 indicates a perfect classifier.

## 3 Results and discussion

### 3.1 Influence of the distance between adjacent residues on the model

The binding sites of proteins typically involve complex interactions with neighboring residues, and changes in the neighboring residues can affect the formation and stability of these interactions, thereby influencing the properties and function of the binding sites (Stank et al., 2016). In this paper, we will use the local neighborhood environment of residues to predict the binding sites of G protein-coupled receptors, where the number of neighboring residues directly affects the richness of structural information contained within the voxel space. The determination of the size of the voxelized space is crucial for deciding the amount of information about the binding site vicinity; the voxel space should not be too large or too small. A space that is too large can cover more protein structure data, but the voxelized structures of residues at the edges may be imbalanced, and too many convolutional layers during the construction of the convolutional network might affect the predictive performance of the 3D CNN to some extent. A space that is too small can avoid overly sparse voxel data but fails to make full use of the structural information of neighboring residues.

Considering that the span of G protein-coupled receptors typically ranges from 40-70Å and the distance between two adjacent amino acids is generally over 3.5Å, we set the edge length of the voxelized three-dimensional space (Sp) to 8-40Å for training and evaluation of the model. The spatial size is kept under 40Å to prevent the voxelized space of a single residue from covering the entire protein structure. Aiming for optimal predictive performance, this section searches for the best size of the voxelized space, using 25% of the training set as the validation set, with epochs set to 150, and employing adam with an adaptive learning rate. Table 2 shows the impact of different sizes of voxelized space on model performance. The results indicate that the size of the voxelized space indeed has an impact on the predictive performance of the model.

**Table 2.** Model performance corresponding to voxel space size.

| Voxelization space(Å<sup>3</sup>) | Accuracy | Precision | Recall | AUC | F1-score | MCC |
| --- | --- | --- | --- | --- | --- | --- |
| Sp=8 | 0.920 | NA | - | 0.5 | NA | - |
| Sp=10 | 0.914 | 0.318 | 0.067 | 0.531 | 0.111 | 0.115 |
| Sp=12 | 0.913 | 0.324 | 0.079 | 0.537 | 0.127 | 0.127 |
| Sp=14 | 0.927 | 0.726 | 0.140 | 0.543 | 0.235 | 0.298 |
| Sp=16 | 0.930 | 0.737 | 0.196 | 0.579 | 0.310 | 0.357 |
| Sp=18 | 0.928 | 0.635 | 0.226 | 0.591 | 0.333 | 0.350 |
| Sp=20 | 0.929 | 0.634 | 0.258 | 0.627 | 0.367 | 0.374 |
| Sp=22 | 0.930 | 0.653 | 0.272 | 0.636 | 0.384 | 0.392 |
| Sp=24 | 0.932 | 0.69 | 0.280 | 0.649 | 0.399 | 0.412 |
| Sp=26 | 0.935 | 0.729 | 0.300 | 0.665 | 0.425 | 0.442 |
| Sp=28 | 0.942 | 0.751 | 0.411 | 0.687 | 0.531 | 0.529 |
| Sp=30 | 0.946 | 0.769 | 0.458 | 0.724 | 0.575 | 0.568 |
| Sp=32 | 0.942 | 0.678 | 0.532 | 0.743 | 0.596 | 0.570 |
| Sp=34 | 0.942 | 0.692 | 0.485 | 0.716 | 0.570 | 0.550 |
| Sp=36 | 0.940 | 0.694 | 0.441 | 0.692 | 0.539 | 0.523 |
| Sp=38 | 0.940 | 0.649 | 0.512 | 0.711 | 0.572 | 0.545 |
| Sp=40 | 0.943 | 0.696 | 0.505 | 0.727 | 0.585 | 0.564 |

According to Table 2, a larger space results in better predictive performance because the input data consists of higher-dimensional tensors, which encompass more comprehensive structural information. When Sp is less than 10Å, the 3D convolutional neural network does not perform effectively in predicting binding sites. The models constructed have fewer convolutional and pooling layers, which are insufficient to extract potential structural relations from the tensor data. Regardless of the size of the space, the accuracy(ACC) of the models is over 90%, due to a larger proportion of negative samples, leading to more precise predictions for labels of 0. Even if predictions for positive samples are inaccurate, the overall level of prediction remains high. However, this metric is not very meaningful for assessing the performance of the model. When Sp equals 32Å, the model achieves the highest F1-score value, indicating superior performance compared to models of other space sizes.

Our dataset is imbalanced, with a smaller number of positive samples. Considering F1-score integrates precision and recall, we use it as the primary criterion for evaluation, with AUC and MCC as secondary conditions. We believe that establishing a convolutional neural network with Sp=32Å enables a better prediction of GPCRs binding sites from structural data. Different input tensors require the construction of different convolutional models, and a series of parameters including the number of convolutional layers, pooling layers, fully connected layers, filter sizes, kernel sizes, and training batches, all affect model performance. It’s not feasible for us to use the control variable method to filter by space size. When building the model, we prioritize increasing the number of convolutional layers and ensure there is at least one pooling layer to reduce the computational complexity of the model. These settings, to a certain extent, reduce the risk of overfitting and aim to ensure the accuracy of our results as much as possible. In our future work, we will use 32Å to establish our input data and explore the prediction of binding sites by convolutional neural networks of different dimensions.

### 3.2 Model performance analysis

Through the deep learning network model established above, the GS108 dataset was trained and predicted. To explore whether 3D convolutional neural networks based on structural prediction of key residues are superior to other models, we also considered 2D convolutional neural networks and 1D convolutional neural networks. For processing the 2D CNN dataset, we projected the voxelized three-dimensional data (32,32,32,4) from the X, Y, Z directions respectively, resulting in (32,32,12) dimensional data which was then used for 2D convolutional neural network training, and comparative analysis with other models. We also compared it with 3D convolutional neural networks not utilizing data enhancement (3D CNNe). To assess the ability of convolutional neural networks to process one-dimensional data, we used the SVM model for comparison, where the 1D CNN input data has the shape of (32768,4). Table 3 summarizes the evaluation metrics for the five models. The results indicate that the 3D CNN model trained on the GS108 dataset scores the highest in AUC, Recall, and F1-score. It surpasses the non-augmented 3D convolutional neural network model (3D CNNe) and the other three machine learning models. For the prediction of key residues, it can achieve a level of 53.2%, which is a notable achievement.

**Table 3.** Performance comparison of different models.

| Model | Accuracy | Precision | Recall | AUC | F1-score | MCC |
| --- | --- | --- | --- | --- | --- | --- |
| 3D CNN | 0.942 | 0.678 | 0.532 | 0.743 | 0.596 | 0.570 |
| 3D CNN<sub>e</sub> | 0.945 | 0.779 | 0.439 | 0.714 | 0.562 | 0.560 |
| 2D CNN | 0.934 | 0.639 | 0.388 | 0.679 | 0.483 | 0.466 |
| 1D CNN | 0.926 | 0.601 | 0.233 | 0.592 | 0.336 | 0.344 |
| SVM | 0.932 | 0.706 | 0.247 | 0.834 | 0.367 | 0.393 |

The dataset has only 9.64% positive samples, yet the model is still able to identify key residues through structure, which indicates that the 3D convolutional neural network is capable of extracting potential correlations from the structural information of proteins. Table 4 compiles the key residues of the protein 5G53 and bolds the residues correctly identified by the model. Figure 4 illustrates the predictive results of the 3D convolutional neural network for the protein 5G53, where green denotes True Positives (TP) and red represents False Positives (FP). Protein 5G53 has a total of 31 key residues, and our model accurately predicted 15 of them while incorrectly predicting 4 residues (Ala15^1.41^, Asp52^2.50^, Phe62^2.58^, Val172^ECL2^). From Figure 4, it’s observable that the model can roughly recognize the binding pocket of GPCRs and effectively differentiate local features from different positions.

**Figure 4.**
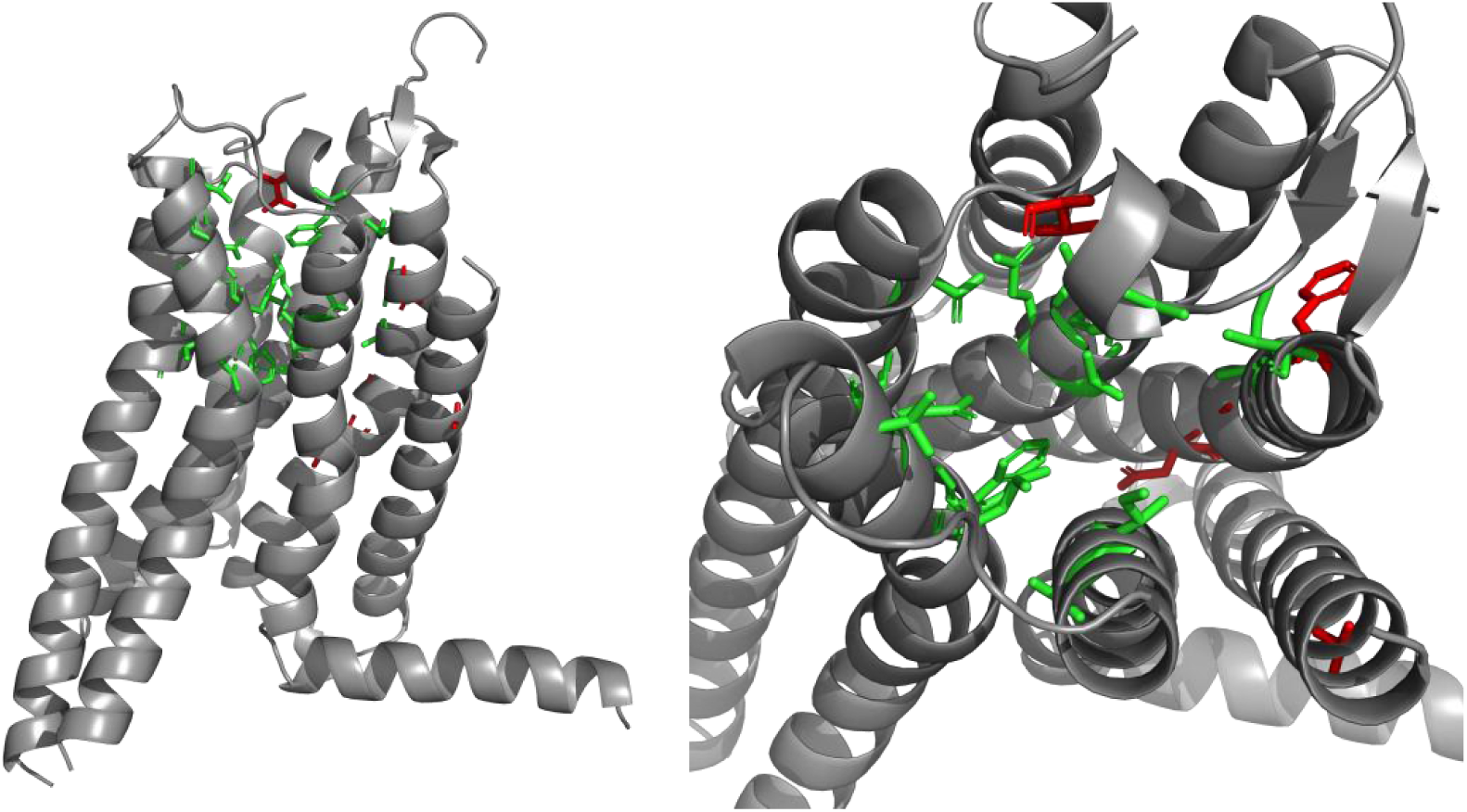
Binding sites predicted by the 3D CNN model.

**Table 4.** The key sites of 3D CNN recognition 5G53 Key residue.

| Key residue |  |  |  |  |  |  |
| --- | --- | --- | --- | --- | --- | --- |
| Tyr9<sup>1.35</sup> | Glu13<sup>1.39</sup> | **Ala59**<sup>2.56</sup> | Ala63<sup>2.60</sup> | **Ile66**<sup>2.63</sup> | Ala81<sup>3.29</sup> | Val84<sup>3.32</sup> |
| **Leu85**<sup>3.33</sup> | **Val86**<sup>3.34</sup> | **Thr88**<sup>3.36</sup> | **Gln89**<sup>3.37</sup> | Ile92<sup>3.40</sup> | **Phe168**<sup>ECL2</sup> | Glu169<sup>ECL2</sup> |
| Met174<sup>5.37</sup> | Met177<sup>5.40</sup> | **Asn181**<sup>5.43</sup> | Cys185<sup>5.46</sup> | **Val186**<sup>5.47</sup> | **Trp246**<sup>6.48</sup> | **Leu249**<sup>6.51</sup> |
| His250<sup>6.52</sup> | Ile252<sup>6.54</sup> | **Asn253**<sup>6.55</sup> | **Thr256**<sup>6.58</sup> | His264<sup>ECL3</sup> | Met270<sup>7.34</sup> | **Ala273**<sup>7.37</sup> |
| **Ile274**<sup>7.38</sup> | Ser277<sup>7.41</sup> | His278<sup>7.42</sup> |  |  |  |  |

Note: The key residues predicted by the model are in bold, and the positive ones are the ligand binding sites confirmed by experiments.

The CNN model established based on one-dimensional structural information also achieved commendable predictive accuracy. It obtained satisfactory results in terms of accuracy (0.926), precision (0.601), and recall (0.233). Although the recall rate is slightly lower than that of SVM and 2D CNN, it can still identify some key residues, but its precision is much lower than anisotropic network models (Su et al., 2016). Our dataset, consisting entirely of G protein-coupled receptors, has a strong homology and very similar structures, which allows the model to have good effects based merely on structure prediction. Since the dataset contains only 108 proteins, obtaining more binding site data of G protein-coupled receptors could better train the model and further improve its accuracy.

### 3.3 Feature Importance Analysis

The impact of input data on classification decision outcomes can be evaluated using gradient information. During the backpropagation algorithm, we can calculate the gradient of the input data, which indicates each element’s contribution to the final classification outcome (LeCun et al., 2015). In this paper, the model is designed with forward and backward propagation processes incorporated, and a loss function is used to compute its loss. Utilizing GradientTape allows us to calculate the gradient of the input data with respect to the loss. By analyzing the gradient values of the original data, we can deduce which local atoms have the most significant impact on the model’s prediction outcome. In this section, the calculated gradients are mapped and dimensionally reduced from three dimensions. Figure 5 shows the mapping of the absolute values of gradients for Thr88^3.36^ from the PDB named 5G53 in three dimensions, and these absolute values of gradients are normalized to facilitate direct observation of important features. The residue Thr88^3.36^ is an important binding site that directly affects the conformational changes during activation and controls the expansion of the “ribosome pocket” (Ciancetta et al., 2015). In the test set, the protein residue structures at this site achieved good predictive results, and the model could generally identify this site as a key residue. Since the central position of the original data contains many atoms, the significance in the central position shown in Figure 5 is also relatively weak, with importance scores mostly maintained at 0.2 and below. However, the importance of the features around the central position is relatively more critical. The structural features at the edges are significantly different, and the model’s gradients are more pronounced there, which fully demonstrates that the variability of the data is one of the key factors in improving model performance. Comparing the input data for the Thr88^3.36^ of different proteins, it is an interesting finding that the greater the local differences in different channels, the more significant the weight of feature importance assigned to the corresponding channel’s structure. Therefore, the model’s predictive capacity can potentially be improved by increasing the number of channels to record more structural information about key residues for model training.

**Figure 5.**
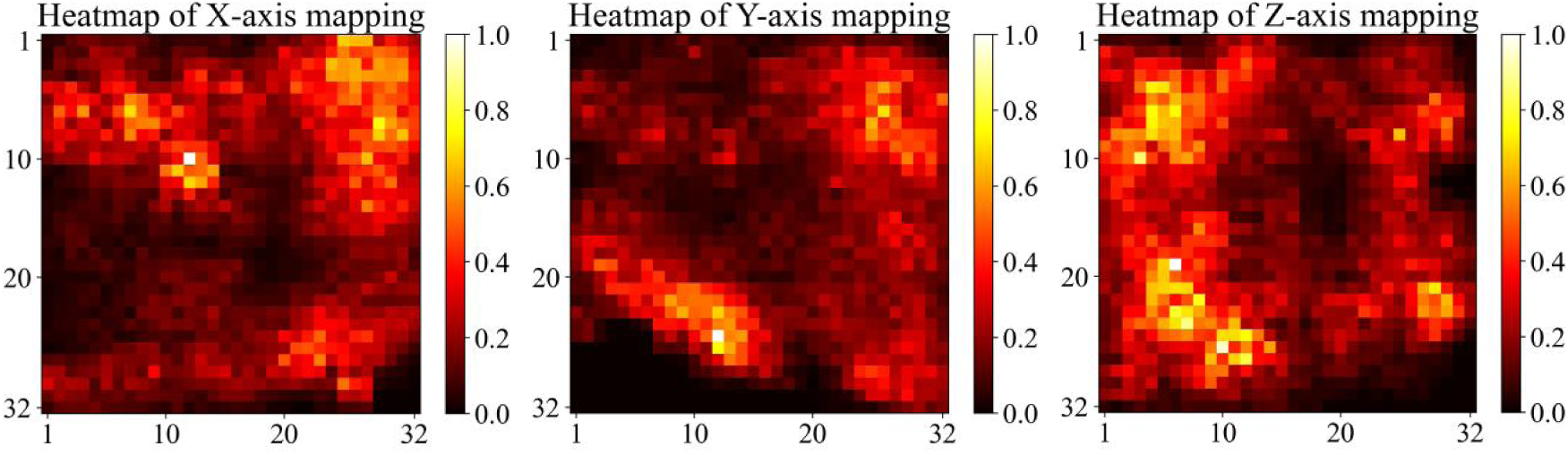
Gradient mapping of different dimensions.

## 4 Conclusion

This research conducted predictive analyses of key residues based on the structure of G protein-coupled receptors. Our approach considered the neighboring structure of amino acids, sectioning off a specific range of amino acids for voxelization in dataset construction, and then establishing a 3D convolutional neural network model. Through an iterative process, we determined the optimal local structure size and demonstrated that key residues could be predicted solely based on the three-dimensional structure of proteins. To compare the predictive accuracy of different models based on structure, we also compared 2D CNN, 1D CNN, SVM, and 3D CNN without data augmentation. The results showed that 3D CNN with data augmentation performed better. To explore which input features impact the determination of key residues, we conducted an analysis of the importance of input data features. We visualized the voxelized protein structure and displayed importance scores through heat map colors, which allowed us to determine the contribution of each element to the final classification decision based on color.

Currently, the scPDB database does not have comprehensive data on G protein-coupled receptor binding sites, which limited our ability to train the model with more data and, to a certain extent, constrained the performance of the deep learning model. Predicting key residues based solely on structural data is also quite limited, as the physicochemical properties of residues are also very important. To improve the prediction accuracy of the model, we should consider an integrated approach that includes protein sequence information, physicochemical properties, and structural information. This work is of significant reference value for research on predicting key protein residues based on structure and aids in the design and discovery of drug molecules.

## Author contributions

All authors contributed to the study conception and design. Material preparation, data collection and analysis were performed by Junfeng Yu. The first draft of the manuscript was written by Junfeng Yu and all authors commented on previous versions of the manuscript. All authors read and approved the final manuscript.

## Supporting information

https://github.com/Yu-0911/GS108dataset.git

## Acknowledgments

The work was supported by the Inner Mongolia Autonomous Region Natural Science Foundation (No. 2022LHMS03014, No. 2024LHMS06018) and Basic Scientific Research Business Fee Project of Colleges and Universities Directly under Inner Mongolia Autonomous Region (JY20220069).

