## Supplementary material for "Prediction of binding sites of GPCRs based on 3D convolutional neural networks": https://github.com/Yu-0911/GS108dataset.git

### Supplementary document

TabA Summary of data information of 108 GPCRs protein

| PDB code | Length | Key residuals | PDB code | Length | Key residuals | PDB code | Length | Key residuals |
|---|---|---|---|---|---|---|---|---|
| 2I35 | 328 | 34 | 4IAQ | 367 | 41 | 5D6L | 442 | 44 |
| 2RH1 | 442 | 40 | 4IAR | 379 | 38 | 5DHG | 281 | 36 |
| 2VT4 | 276 | 30 | 4IB4 | 375 | 42 | 5DHH | 282 | 33 |
| 2X72 | 326 | 32 | 4JKV | 454 | 45 | 5DSG | 399 | 31 |
| 2Y00 | 298 | 31 | 4K5Y | 248 | 33 | 5F8U | 276 | 30 |
| 2Y01 | 286 | 54 | 4LDE | 454 | 35 | 5JQH | 391 | 30 |
| 2Y02 | 298 | 58 | 4LDL | 454 | 32 | 5L7D | 578 | 26 |
| 2Y03 | 298 | 51 | 4MBS | 346 | 39 | 5L7I | 572 | 36 |
| 2Y04 | 298 | 52 | 4MQT | 275 | 27 | 5TGZ | 439 | 34 |
| 2YCW | 297 | 26 | 4N4W | 457 | 42 | 5XR8 | 431 | 23 |
| 2YCX | 286 | 29 | 4N6H | 408 | 30 | 5XRA | 428 | 57 |
| 2YCY | 288 | 33 | 4NC3 | 370 | 41 | 2YDO | 309 | 27 |
| 2YCZ | 291 | 29 | 4NTJ | 369 | 35 | 2YDV | 315 | 32 |
| 3D4S | 439 | 30 | 4O9R | 441 | 32 | 3EML | 448 | 24 |
| 3NY8 | 439 | 28 | 4OR2 | 366 | 10 | 3PWH | 291 | 29 |
| 3NY9 | 439 | 31 | 4PHU | 434 | 35 | 3QAK | 444 | 41 |
| 3NYA | 439 | 29 | 4PXZ | 392 | 43 | 3REY | 291 | 31 |
| 3ODU | 466 | 27 | 4PY0 | 389 | 43 | 3RFM | 291 | 20 |
| 3OE9 | 420 | 25 | 4QIM | 457 | 41 | 3UZA | 291 | 28 |
| 3P0G | 284 | 34 | 4QIN | 447 | 39 | 3UZC | 291 | 26 |
| 3PBL | 432 | 30 | 4S0V | 478 | 35 | 3VG9 | 297 | 26 |
| 3SN6 | 443 | 34 | 4U14 | 436 | 29 | 3VGA | 296 | 23 |
| 3UON | 438 | 31 | 4U15 | 392 | 32 | 4EIY | 390 | 26 |
| 3V2W | 442 | 32 | 4U16 | 392 | 26 | 4UG2 | 292 | 39 |
| 3V2Y | 455 | 32 | 4XNW | 346 | 34 | 4UHR | 310 | 37 |
| 3VW7 | 442 | 37 | 4Z34 | 385 | 40 | 5G53 | 283 | 31 |
| 3ZPQ | 299 | 47 | 4Z35 | 382 | 43 | 5IU4 | 388 | 25 |
| 3ZPR | 298 | 32 | 4Z36 | 382 | 40 | 5IU8 | 387 | 24 |
| 4AMI | 282 | 30 | 4Z9G | 421 | 33 | 5IUA | 388 | 28 |
| 4AMJ | 299 | 33 | 4ZJ8 | 503 | 36 | 5IUB | 388 | 28 |
| 4BVN | 289 | 31 | 4ZJC | 503 | 35 | 5JTB | 388 | 24 |
| 4DAJ | 429 | 29 | 5A8E | 285 | 32 | 5K2A | 398 | 25 |
| 4DJH | 447 | 37 | 5C1M | 295 | 33 | 5K2B | 394 | 26 |
| 4EA3 | 278 | 33 | 5CXV | 444 | 33 | 5K2C | 396 | 26 |
| 4EJ4 | 442 | 26 | 5D5A | 442 | 27 | 5K2D | 395 | 26 |
| 4GBR | 286 | 26 | 5D5B | 442 | 40 | 5UVI | 391 | 27 |
